# The crosstalk between microbial sensors ELMO1 and NOD2 shape intestinal immune responses

**DOI:** 10.1101/2022.07.09.499433

**Authors:** Aditi Sharma, Sajan Chandrangadhan Achi, Stella-Rita Ibeawuchi, Mahitha Shree Anandachar, Hobie Gementera, Uddeep Chaudhury, Fatima Usmani, Kevin Vega, Ibrahim M Sayed, Soumita Das

## Abstract

Microbial sensors play an essential role in maintaining cellular homeostasis. Our knowledge is limited on how microbial sensing helps in differential immune response and its link to inflammatory diseases. Recently, we have shown that cytosolic sensor ELMO1 (Engulfment and Cell Motility Protein-1) binds to effectors from pathogenic bacteria and controls intestinal inflammation. Here, we show that ELMO1 interacts with another sensor, NOD2 (Nucleotide-binding oligomerization domain-containing protein 2), that recognizes bacterial cell wall component muramyl dipeptide (MDP). The polymorphism of NOD2 is linked to Crohn’s disease (CD) pathogenesis. Interestingly, we found that overexpression of ELMO1 and mutant NOD2 (L1007fs) were not able to clear the CD-associated adherent invasive *E. coli (AIEC*-LF82). To understand the interplay of microbial sensing of ELMO1-NOD2 in epithelial cells and macrophages, we used enteroid-derived monolayers (EDMs) from ELMO1 and NOD2 KO mice and ELMO1 and NOD2-depleted murine macrophage cell lines. The infection of murine EDMs with *AIEC*-LF82 showed higher bacterial load in ELMO1-KO, NOD2 KO EDMs, and ELMO1 KO EDMs treated with NOD2 inhibitors. The murine macrophage cells showed that the downregulation of ELMO1 and NOD2 is associated with impaired bacterial clearance that is linked to reduced pro-inflammatory cytokines and reactive oxygen species. Our results indicated that the crosstalk between microbial sensors in enteric infection and inflammatory diseases impacts the fate of the bacterial load and disease pathogenesis.

## INTRODUCTION

The intestinal epithelium has enormous surface area and harbors a plethora of microbes which includes bacteria, fungi, viruses and protozoa ^1^. Although these microbes are separated by biophysical and biochemical barriers, they are in constant interaction with the epithelial cells and host immune system. Any changes in the composition of the microbes lead to perturbations in the barrier and can disrupt intestinal homeostasis. Disruption of intestinal homeostasis has been accounted for various chronic inflammatory diseases such as inflammatory bowel diseases, cardiovascular diseases, metabolic disorders and cancer ^1,2^. Intestinal epithelium plays a critical role in maintenance of homeostasis. They are involved in constant sampling of the intestinal microenvironment, sensing of commensals and pathogens, secretion of compounds, and triggering immune response which influences the colonization of the microbes ^3,4^.

Pattern recognition receptors (PRRs) are involved in sensing microbes and can distinguish commensals and pathogens by identifying pathogen-associated molecular patterns associated with microbes ^5^. One such protein is EnguLfment and cell MOtility protein 1 (ELMO1), which has been recently implicated in having a role in microbial sensing. ELMO1 facilitates bacterial internalization, mounts inflammatory response, and coordinates bacterial clearance ^6–9^. Brain Angiogenesis Inhibitor-1 (BAI1) recognizes bacterial lipopolysaccharide, interacts with ELMO1 resulting in the activation of Rac1 and induction of pro-inflammatory cytokines ^10^. Studies from our group have revealed that ELMO1 is involved in the sensing of microbes associated with IBD ^11^. Using enteroid-derived monolayers (EDMs) from the organoids isolated from colonic biopsies of IBD patients, we have demonstrated that bacterial internalization and production of pro-inflammatory cytokines such as TNF-α and MCP-1 are ELMO1-dependent, which consequently triggers the chronic inflammatory cascade of IBD ^11^. Interestingly, we found that ELMO1 induces differential immune responses between pathogens and commensals by interacting with several bacterial effectors containing the WxxxE motif ^9^

NOD2 is a cytosolic PRR belonging to the Nod-like receptor (NLR) family that recognizes processed bacterial cell wall component, muramyl dipeptide (MDP), from both Gram-positive and Gram-negative bacteria ^12^. NOD2 activation results in recruitment of RICK (RIP-like interacting CLARP kinase)/RIP2 (Receptor-Interacting Protein 2) and activation of nuclear factor-κB (NF-κB) and mitogen-activated protein kinase (MAPK) pathways. NOD2 is also involved in secretion of cytokines such as IL-8, upregulation of pro-interleukin 1β, induction of autophagy, production of antimicrobial peptides and defensins, and maintenance of intestinal homeostasis ^12–15^. NOD2 plays a crucial role in regulation of microbial flora in intestine ^16,17^. NOD2 deficient mice had a reduced capability to prevent colonization of pathogenic microbes in the intestine and had impaired bactericidal activity ^18^. NOD2 has been identified to have a significant role in inflammatory bowel disease especially CD. Mutations in *NOD2* and variants of *NOD2* have been shown to increase susceptibility to CD ^19,20^. Single nucleotide polymorphisms (SNPs) and variation in NOD2 receptors were recorded in CD patients, specifically two missense mutations, R702W and G908R, and one frameshift mutation, L1007fs ^19,20^. NOD2 variants associated with CD has been shown to be defective in the recognition of MDP ^21^. Bacterial clearance is impaired in CD, wherein the mutations of *NOD2* associated with CD are accompanied by impaired initiation of autophagy and bacterial elimination. However, the mechanisms through which *NOD2* mutations lead to enhanced inflammation are not completely understood.

ELMO1 and NOD2 are cytosolic sensors and are also implicated in IBD, however whether these sensors interact with each other and the influence of such interaction on bacterial pathogenesis is not known. In this study, we assessed if NOD2 and ELMO1 could interact with each other directly or indirectly to regulate the bacterial sensing. We used stem-cell based approaches to recapitulate normal gut physiology, and intestinal bacteria, *AIEC*-LF82, as stressor to assess gut function. We found that both these cytosolic proteins interact with each other and regulate bacterial load, reactive oxygen species (ROS) generation and induction of pro-inflammatory cytokine response in macrophages and gut epithelial cells. Overall, we found that the interactions of these two bacterial sensors are not only important for bacterial sensing but also regulate the outcome of inflammation during enteric infection and inflammatory diseases.

## RESULTS

### ELMO1 interacts with cytosolic microbial sensor NOD2

Previously, we have demonstrated that ELMO1 is involved in microbial sensing and control of intestinal inflammation ^6^. Both ELMO1 and NOD2 are bacterial sensor proteins that play a role in autophagy and facilitate bacterial degradation ^22,23^. Since ELMO1 and NOD2 regulate intestinal immune response against bacterial infections ^6–8,16,24–27^, we assessed if ELMO1 and NOD2 could interact physiologically. We transiently expressed HA-tagged NOD2, and Flag tagged ELMO1 in HEK293 cells and performed co-immunoprecipitation. Immunoprecipitation of ELMO1 with FLAG antibody showed an interaction of ELMO1 and NOD2 (**Figure 1A**). To determine the interacting domains between ELMO1 and NOD2, GST-ELMO1 full length (FL) and GST-ELMO1 C-terminal domain (CT) were immobilized on glutathione beads and GST pulldown assay was performed with recombinant His-NOD2 CARD or His-NOD2 LRR. Our data suggests that only NOD2 LRR region could bind ELMO1 and that the CT domain of ELMO1 is sufficient for this interaction (**Figure 1B**). GST pulldown assay using GST NOD2 LRR with His-ELMO1 FL or His-ELMO1 CT, showed that the CT domain of ELMO1 had higher binding compared to the FL ELMO1 (**Figure 1C**). Similar findings were observed when recombinant His-ELMO1-CT and His-ELMO1-FL was used in GST pulldown assays with different concentrations of GST or GST-NOD2-LRR (**Figure S1**).

**Figure 1.**
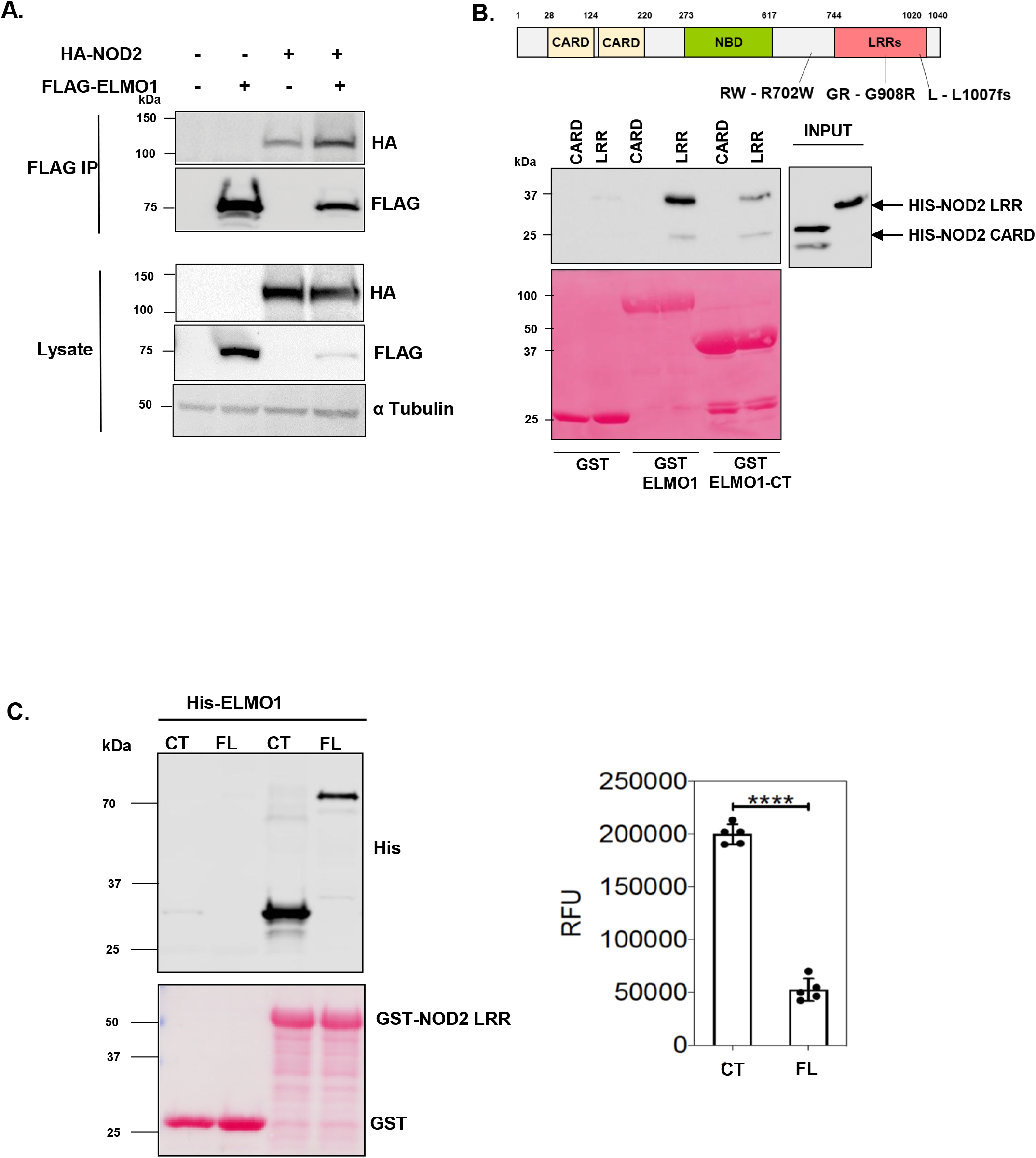
The C terminal part of ELMO1 interacts with the LRR domain of NOD2. **A.** HEK293 cells were co-transfected with Flag-ELMO1 and HA-NOD2. After transfections, cells were lysed, normalized for protein content and precipitated using anti-FLAG antibody. Immunoprecipitants and cell lysates were visualized by immunoblotting with corresponding antibodies. **B.** The schematic shows the structure of NOD2 protein and associated mutations involved in Inflammatory Bowel Disease (IBD). To detect the regions of NOD2 that bind ELMO1, Pulldown assay using GST, GST-ELMO1 full length (FL) and GST-ELMO1 C-terminal (CT) were immobilized on glutathione beads. The soluble recombinant His-NOD2 CARD or His-NOD2 LRR proteins were incubated with the beads. Bound NOD2 proteins in the pull down and in the input were determined using anti-His antibody. The ponceau staining in the lower panel showed the equal loading of GST tagged proteins. **C.** Recombinant His-ELMO1 FL and His-ELMO1 CT were used in GST pulldown assays with GST or GST-NOD2-LRR. Bound His-ELMO1 FL or His-ELMO1 CT were visualized by immunoblotting using anti-His antibody. The ponceau staining in the lower panel showed the equal loading of GST tagged proteins. The densitometry of the pulldowns was shown in the graph from three independent experiments where the t-test showed the p value of < 0.0001 as ****.

The LRR domain is required for MDP binding that unfolds NOD2 from auto-inhibitory state to its active state ^28,29^. Since most implicated mutations in NOD2 are present within and around the LRR domain, we next evaluated the interaction of ELMO1 with selected NOD2 mutants (GR - G908R, RW - R702W, L - L1007fs), that are associated with susceptibility to CD ^19,20^. HEK293 cells were transiently transfected with FLAG-ELMO1 and HA-NOD2 WT (Wild Type) or HA-NOD2 mutants (GR - G908R, RW - R702W, L - L1007fs) followed by co-immunoprecipitation with FLAG antibody. Our results showed that there were no significant differences in the binding of NOD2 mutants with ELMO1 when compared to NOD2 WT (**Figure 2A**). This data indicated that these CD-associated NOD2 mutations did not alter the interaction of NOD2 with ELMO1.

**Figure 2.**
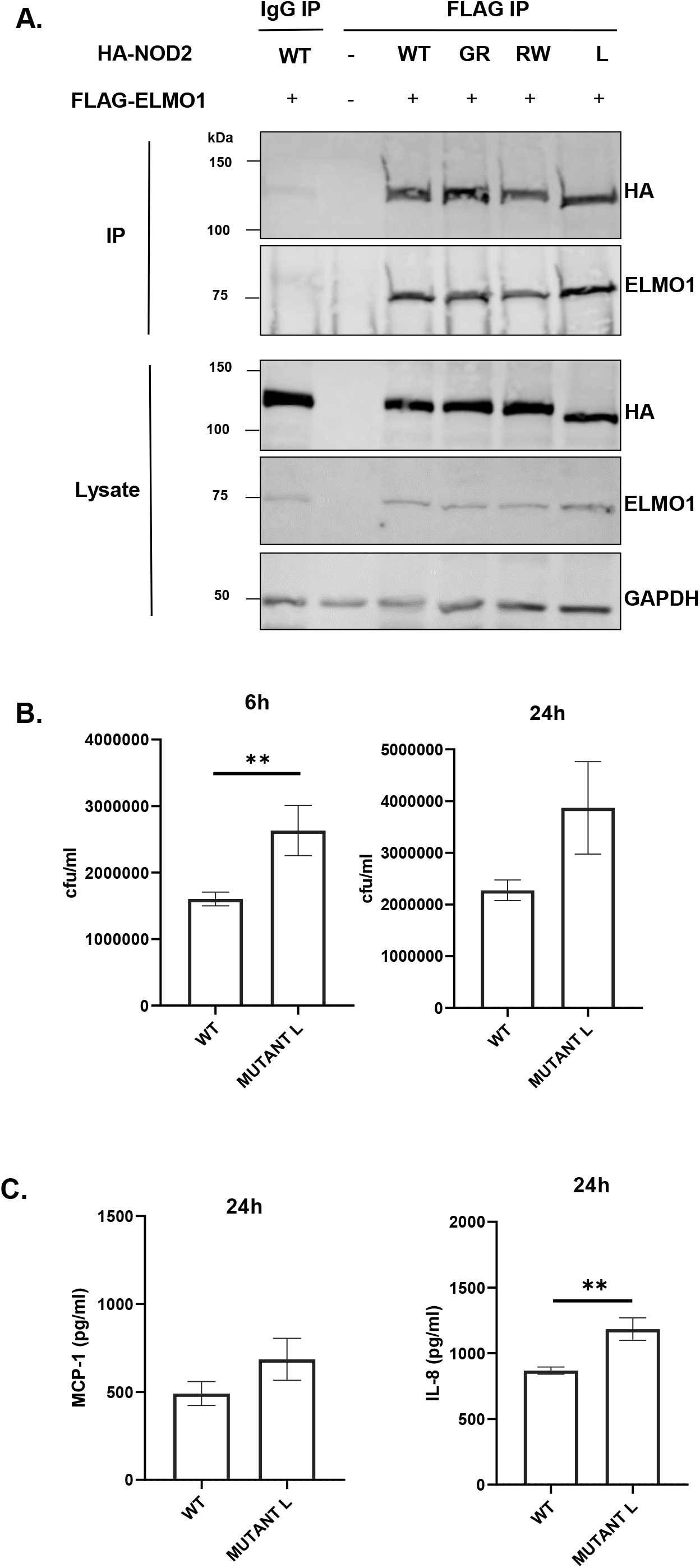
The involvement of NOD2 mutant in bacterial clearance and in inflammation. **A.** HEK293 cells were co-transfected with Flag-ELMO1 and with HA-NOD2 [WT and mutant (GR - G908R, RW - R702W, L - L1007fs)]. After transfections, cells were lysed, normalized for protein content and precipitated using anti-FLAG antibody. Immunoprecipitants and cell lysates were visualized by immunoblotting with corresponding antibodies. **B.** HEK293 cells were transfected with vectors over-expressing ELMO1 and BAI1, with either NOD2 WT or NOD2L1007fs (Mutant L). Cells were infected with AIEC-LF82 for 6h and gentamicin was used to kill extracellular bacteria. The bacterial count at 6 h and 24 h were plotted from 3 different experiments. **C.** The supernatant from B at 24 h was used to measure cytokine by ELISA. Data represented as mean ± SEM. Statistical significance was estimated using Mann Whitney test where **p < 0.01.

To further understand the physiological relevance of this interaction, we co-transfected HEK293 cells with wild type NOD2 plasmid or NOD2 L1007fsinsC (L1007fs or mutant L) mutant in the presence of ELMO1 and BAI1. We used the L1007fs mutant as this frameshift mutation is most common in CD patients. We found higher bacterial load of *AIEC*-LF82 infection in mutant L compared to WT at two different time points (6 h and 24 h post infection) (**Figure 2B**). Interestingly, the mutant L was also associated with higher levels of pro-inflammatory cytokines - IL-8 and MCP-1 after 24 h of infection with *AIEC*-LF82 (**Figure 2C**). Pathogens are known to reduce host NF-kB activity in order to enhance their entry by decreasing pro-inflammatory cytokines ^30,31^. NFkB activity measured by luciferase reporter assay showed lower NF-kB activity in the mutant L compared to WT (**Figure S2A**). MDP is the ligand for NOD2 activation, which leads to IL-8 production, so when HEK293 cells were treated with MDP for 6 h, reduced levels of IL-8 were observed in the mutant L as compared to WT (**Figure S2B**). Collectively, our data showed that mutation in NOD2 does not affect its interaction with ELMO1 however it was associated with increased bacterial burden and modulated immune response.

### ELMO1-NOD2 interaction fine-tunes paracellular permeability and bacterial load in the epithelium

An intact epithelial barrier spatially segregates luminal microbes and protects the host from invasion and dissemination of these microbes. There is substantial evidence that NOD2 regulates intestinal barrier function through myosin light chain kinase (MLCK) activity and mutations in NOD2 causes barrier defects in mice ^32^. To assess the impact of NOD2 on the integrity of gut barrier following infection, we challenged EDMs from WT, ELMO1 KO and NOD2 KO mice with *AIEC*-LF82 infection and then assessed the gut barrier integrity by measuring the transepithelial electrical resistance (TEER). We found that NOD2 KO cells had higher paracellular permeability and low resistance to flow-through bacteria on apical side compared to WT cells after 8 h of *AIEC*-LF82 infection (**Figure 3A**). There was no significant difference in TEER values of ELMO1 deficient cells compared to WT cells. Next, to inhibit NOD2 signaling in ELMO1 KO EDMs, we used a potent small molecule NOD2 inhibitor, GSK717. Interestingly, ELMO1 KO EDMs treated with GSK717 showed reduced paracellular permeability with high resistance compared to NOD2 KO EDMs (**Figure 3A**) suggesting that NOD2 is essential for gut barrier integrity.

**Figure 3.**
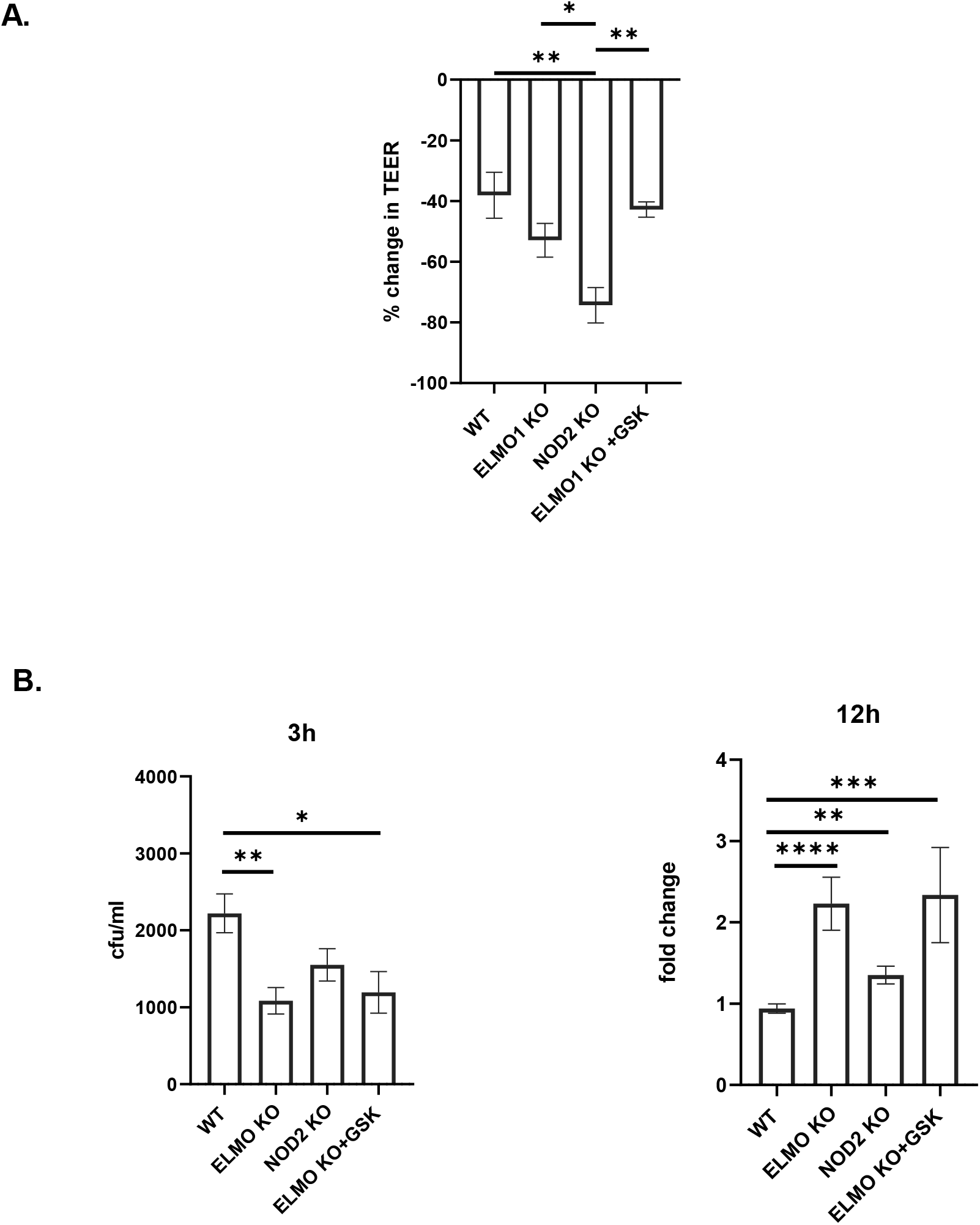
The impact of ELMO1 and NOD2 in bacterial entry and viability in murine ileal EDMs. **A.** Enteroid-derived monolayers (EDMs) from age and gender matched WT, NOD2 KO and ELMO1 KO mouse were infected with *AIEC*-LF82 for 6h. GSK717, a pharmacologic inhibitor of NOD2 was used in ELMO1 KO EDMs to inhibit NOD2 signaling. Transepithelial electrical resistance (TEER) was measured at time intervals and plotted as percentage change at 6h compared to 0h. **B.** The same EDMs from A were infected with *AIEC*-LF82 for 3h. Extracellular bacteria were killed with gentamicin and bacterial load was measured at 12h as described in the materials and methods. The no of internalized bacteria at 3h was plotted as colony-forming units (cfu/mL) and bacterial survival was measured at 12h where the fold change was calculated considering WT EDMs as 1.

We previously showed that ELMO1 regulates bacterial entry in intestinal cells, so we further assessed the impact of interaction on ELMO1 and NOD2 in bacterial internalization and survival in intestinal epithelium. We infected murine ileum EDMs with *AIEC-*LF82 and assessed bacterial entry and survival. Since mouse EDMs lack CEACAM receptors needed specifically by *AIEC*-LF82 for invasion into intestinal cells, the number of internalized bacteria could be low (**Figure 3B**). Like our previous study ^11^ and as shown in **Figure 3B**, ELMO1 KO ileum EDMs had lower number of internalized bacteria after 3h of infection, as compared to both WT and NOD2 KO ileum EDMs. Although a smaller number of bacteria entered in ELMO1 KO cells, the bacterial count was higher at 12h of infection as compared to WT ileum EDMs, depicting prolonged survival and delayed clearance. Similar to ELMO1 deficient EDMs, NOD2 KO EDMs and ELMO1 KO EDMs treated with GSK717 also had a defect in bacterial internalization, and a delayed clearance of bacteria thus leading to higher load after 12 h of infection (**Figure 3B**).

We have shown previously that ELMO1 present in phagosome regulates bacterial clearance in macrophages ^13,23^. Further, ELMO1 KO mice showed reduced colonic inflammation and pro-inflammatory cytokines following intestinal pathogen infection ^9,26^. Here, we isolated bone marrow-derived macrophages (BMDM) from NOD2 KO mice to validate our findings in EDMs. BMDMs from WT and NOD2 KO mice were infected with LF82 or treated with NOD2 ligand MDP (**Figure S3**). We found the secretion of pro-inflammatory cytokine KC and TNF in BMDMs were low in NOD2 KO mice compared to WT when infected with LF82 or treated with MDP. (**Figure S3A-C**). Also, the bacterial burden in BMDMs from NOD2 KO mice was significantly higher compared to WT (**Figure S3D**).

### ELMO1-NOD2 interaction in macrophages regulates bacterial survival, immune responses, and ROS in macrophages

To investigate the crosstalk between ELMO1 and NOD2 in macrophages, we used lentiviral vectors expressing shRNA to knockdown either ELMO1(E1), NOD2(N2) or both (E1N2) in murine macrophages and, compared them to macrophages with control shRNA (C1). As depicted in **Figure S4A**, NOD2 shRNA resulted in knockdown of *NOD2* transcript in both N2 and E1N2 cells. The downregulation of ELMO1 in E1 and E1N2 cells were confirmed by western blot (**Figure S4B**). Bacterial sensing is critical in the generation of immune response. Since both NOD2 and ELMO1 are bacterial sensors, we further investigated the effect of their knockdown on bacterial, survival and induction of innate immune response in macrophages. We infected C1, E1, N2 and E1N2 macrophages with *AIEC-*LF82 and evaluated bacterial burden. We observed that the absence of either or both of the proteins resulted in delayed bacterial clearance which led to higher bacterial load compared to control cells (**Figure 4A**). Previously, we had reported that ELMO1 regulated the immune response against infection in macrophages by producing pro-inflammatory cytokines ^9,26^. In the present study, we evaluated the role of both ELMO1 and NOD2 in the induction of innate immune responses by measuring the levels of pro-inflammatory cytokines in macrophages upon infection with CD-associated *AIEC*-LF82. In *AIEC*-LF82 infection, the absence of either ELMO1 or NOD2 or both resulted in a significant decline in IL-6 levels compared to C1 cells (**Figure 4B**).

**Figure 4.**
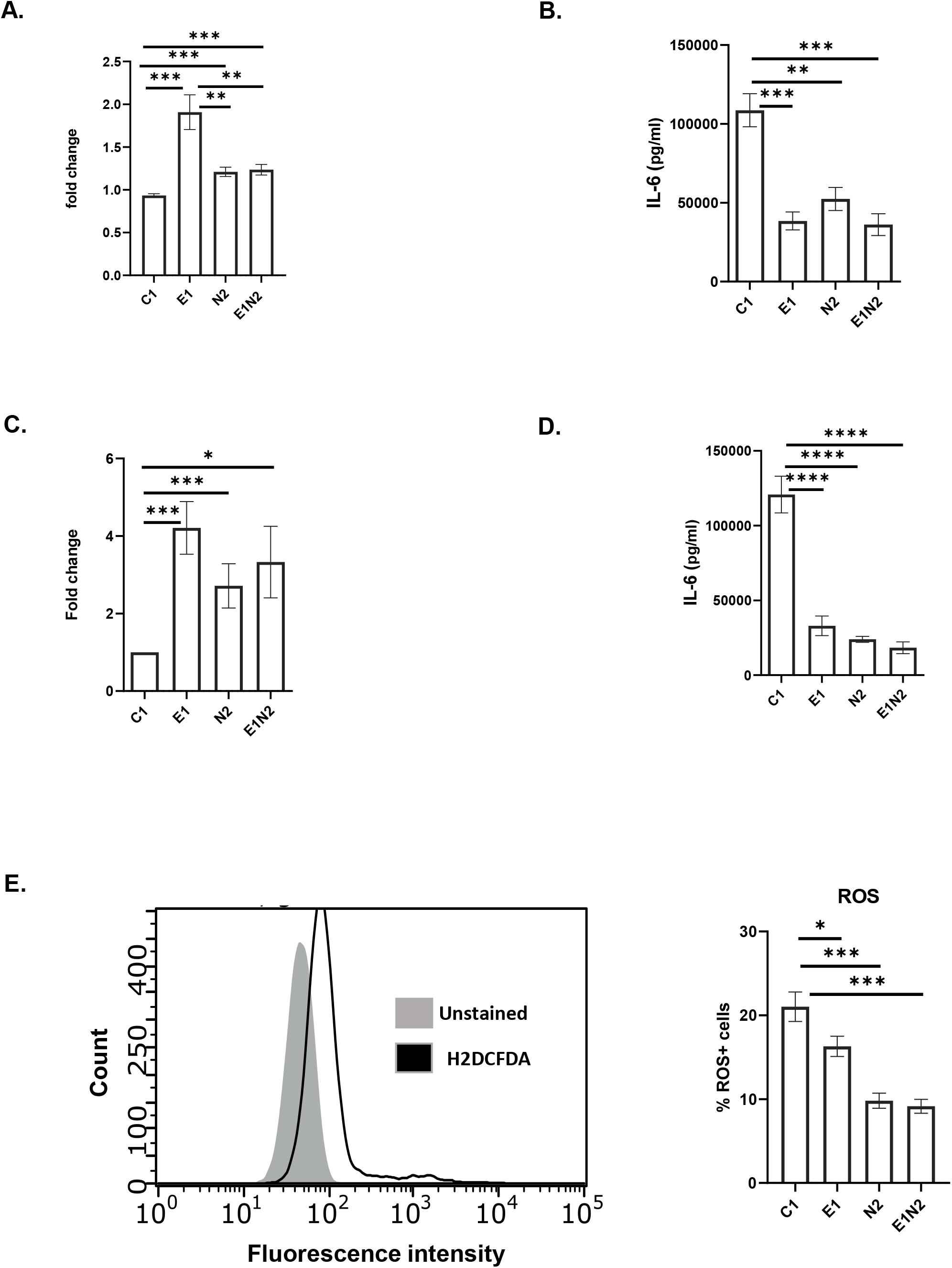
The impact of ELMO1-NOD2 interaction in bacterial clearance and inflammation in macrophages after AIEC-LF82 infection. **A.** J774 cells were infected with *AIEC*-LF82 for 1h, after which extracellular bacteria were killed by gentamicin treatment. Cells were incubated for 12h, then lysed, serially diluted, and plated for colony-forming units (cfu). For bacterial survival at 12h, the cfu at 12h were normalized to the cfu of bacterial entry at 1h for each of the respective macrophages. The graph represents the fold change in the bacterial survival, at 12h calculated by considering bacterial survival of C1 as 1 and the relative bacterial survival in other cells are compared to C1. **B.** Bar graphs display the concentration of pro-inflammatory cytokines (IL-6) released in supernatant collected at end of 12h from ‘A’. **C.** J774 cells were infected with *Salmonella enterica* serovar Typhimurium strain SL1344 for 1h, after which extracellular bacteria were killed by gentamicin treatment. Cells were incubated for 12h, then lysed, serially diluted, and plated for cfu. For bacterial survival at 12h, the cfu at 12h were normalized to the cfu of bacterial entry at 30 min for each of the respective macrophages. The graph represents the fold change in the bacterial survival, at 12h calculated by considering bacterial survival of C1 as 1 and the relative bacterial survival in other cells are compared to C1. **D.** Bar graphs display the concentration of pro-inflammatory cytokines (IL-6) released in supernatant collected at end of 12h from ‘C’. **E.** J774 macrophages were infected with *AIEC*-LF82 infection for 30 min, followed by treatment with high gentamicin and 1uM H2DCFDA for 60 min. Cells were then washed and analyzed on a flow cytometer for detection of total cellular ROS. Bar graph on right show percent of cells expressing ROS, data is displayed as mean ± SEM. Graph shows transcript levels after infection. Data is displayed as mean ± SEM. Statistical significance was estimated using Mann Whitney test and one-way Anova *p < 0.05, **p < 0.01 and ***p < 0.001.

We also validated the above results using *Salmonella enterica* serovar Typhimurium strain SL1344 as a model for enteric infection. As observed for *AIEC*-LF82, in case of *Salmonella* infection the absence of either ELMO1 or NOD2 or both resulted in a delayed bacterial clearance (**Figure 4C**) and significant decline in levels of IL-6 (**Figure 4D**) and IL-1β (**Figure S5A**) compared to C1 cells. Taken together, these results showed that ELMO1 and NOD2 are required for the clearance of *AIEC*-LF82 and *Salmonella* and induction of innate immune response in the immune cells.

ROS production is a natural anti-bacterial response in macrophages against almost all microbes, which reduces the bacterial survival ^33^. But some microbes modulate this effect in order to survive inside the host cells. Since we observed higher bacterial survival in absence of ELMO1 or NOD2 or both, we assessed the ROS levels in these cells. We found lower ROS production in E1, N2 and E1N2 cells compared to C1 cells, which shows that the anti-immune response in macrophages was hijacked by microbes in absence of bacterial sensors (**Figure 4E**). To further confirm this, we isolated peritoneal macrophages from WT, Het and ELMO1 KO mice and infected with LF82 followed by measurement of ROS. Again, we found lower ROS levels in ELMO1 KO mouse as compared to WT or Het mouse (**Figure S5B**). To confirm if high bacterial load is a consequence of low ROS release in macrophages (C1, E1, N2, E1N2), we used ROS scavenger; N-acetyl cysteine (NAC) which reduces ROS levels. We found higher bacterial survival in *AIEC*-LF82 infected cells treated with NAC compared to untreated cells (**Figure S5C**). However, there was no significant difference in bacterial levels among ELMO1 KO and NOD2 KO cells upon addition of NAC as the ROS level in these cells were already very low. This result indicated that lower ROS levels in ELMO1 and NOD2 deficient cells are responsible for delayed bacterial clearance.

## Discussion

Bacterial sensing is considered to be the rate-limiting step in combating any infection. Impaired bacterial sensing has been implicated as the cause of several auto-immune and inflammatory diseases, including CD ^34,35^. In the present study, we have reported for the first time that there is direct interaction of two important microbial sensing proteins, NOD2 and ELMO1, which plays a significant role in determining host response to pathogens. The salient features of the study are: 1. C terminal region of ELMO1 is sufficient for interaction with NOD2; 2. LRR region of NOD2 is involved in binding to ELMO1; 3. The absence of either or both the proteins results in dysregulated antibacterial response.

The C terminal region of ELMO1 is involved in the interaction of DOCK 180 through its PH domain, resulting in the regulation of Rac, resulting in bacterial engulfment and immune response ^36^. In addition, the C-terminal of ELMO1 is involved in binding with bacterial effectors and plays a role in bacterial internalization, dissemination, and induction of host immune responses ^9^. Previously we had shown that BAI1 acts as PRR and identifies the core component of LPS ^37^. The LRR region of NOD2 is similarly involved in the recognition of bacterial components and has been identified to bind to bacterial cell wall component MDP ^12^. In addition, mutations in NOD2 are associated with delayed bacterial clearance in CD patients ^34^. Therefore, the interaction between ELMO1 and LRR region of NOD2 implies coordination between two different bacterial sensing systems, which could affect bacterial recognition, bacterial engulfment and clearance, and regulation of the immune response. Previously, we have reported that the cross-talk between the gut epithelium and immune cells and the axis of epithelial MCP-1 / macrophage TNF-α in the presence of ELMO1 affect the pathogenesis of IBD and bacterial engulfment ^11^. Herein, we studied the effect of ELMO1-NOD2 interaction on bacterial pathogenesis in both epithelial and immune cells.

Mutations in the LRR region of NOD2 have been associated with CD ^19,20^. In this study, we found that although the mutations do not affect the interaction between ELMO1 and NOD2, they affect the clearance of bacteria. Our studies using HEK293 cell lines and mutant NOD2 have substantiated that the interaction of ELMO1-NOD2 does not affect the internalization of bacteria; however, the subsequent immune signaling is dysregulated. L1007fsinsC mutation has been associated with CD and results in the production of truncated NOD2 protein. L1007fsinsC NOD2 mutant has been previously reported to decrease the NFkB activity compared to wild-type NOD2 ^19^. As shown in the previous study ^19^, our study also confirmed the lower NFkB activity in L1007fsinsC mutant, which probably resulted in impaired clearance of bacteria.

*In vivo* and *in vitro* studies have demonstrated the role of NOD2 in maintaining permeability in epithelial cells ^38,39^. In a parallel line, our study using EDM models has shown that the absence of NOD2 decreases the barrier integrity resulting in a higher drop in transepithelial electrical resistance. However, a similar effect was not observed in case of ELMO KO and GSK treated cells suggesting that only NOD2 is vital for gut barrier integrity.

Both ELMO1 and NOD2 are involved in bacterial clearance. ELMO1 plays a role in the engulfment, pathogenesis, and immune responses against enteric bacteria ^9,26^. ELMO1 has also been associated with LC3 associated phagocytosis, induction of inflammatory cytokines and clearance of bacteria and defects in ELMO1 expression results in reduced clearance of bacteria ^23^. NOD2 has been reported to play a role in autophagy, ROS generation, regulation of cytokines and itself can act as antibacterial agent ^12–15^. Defects in NOD2 expression or absence of NOD2 can hence result in delayed clearance of microbes ^34^. Herein, we assessed if the interaction between ELMO1 and NOD2 could affect the clearance of bacteria in both epithelium and macrophage levels. Epithelial monolayers lacking either or both proteins resulted in higher bacterial load (i.e. delayed clearance). Similarly, higher bacterial loads were recorded in macrophages depleted with either ELMO1 or NOD2 or both compared to control macrophage cells. Collectively, these findings showed that ELMO1, NOD2, and their interaction are important in bacterial clearance and pathogenesis.

To investigate the mechanisms of bacterial survival in these cells, we assessed inflammatory cytokines and ROS levels in these cells. As expected, depletion of ELMO1 and/or NOD2 resulted in diminished inflammatory immune responses as shown by reduced level of inflammatory cytokines following the challenge with enteric pathogens. In a parallel line, the level of ROS was reduced in absence of either protein, but the decrease in ROS level was more prominent when NOD2 is depleted (**Figure 4E**).

In conclusion: our study demonstrates the potential role of ELMO1-NOD2 interaction in the pathogenesis of enteric bacteria by affecting bacterial survival/clearance and regulation of immune response and ROS production. Further studies are required to explore the nature of this interaction and the subsequent pathways involved. Such studies will provide alternative target for therapeutics in case of chronic inflammatory diseases such as CD wherein defective sensing of luminal bacteria in predisposed genetic background has been speculated as the major cause.

## Materials and methods

### Ethics statement

All methods involving human and animal subjects were performed in accordance with the relevant guidelines and regulations of the University of California San Diego and the NIH research guidelines.

### Animals

WT, ELMO1^-/-^, and NOD2^-/-^ C57BL/6 mice were gender- and age-matched littermates which were used to isolate intestinal crypts. NOD2^-/-^ mice were purchased from Jackson Laboratories. Animals were bred, housed, used for all the experiments and euthanized according to the University of California San Diego Institutional Animal Care and Use Committee (IACUC) policies under the animal protocol number S18086. All methods were carried out in accordance with relevant guidelines and regulations and the experimental protocols were approved by institutional policies and reviewed by the licensing committee.

### Bacterial strains and growth conditions

Adherent Invasive *Escherichia coli* strain LF82 (*AIEC-*LF82), isolated from the specimens of Crohn’s disease patient, was obtained from Arlette Darfeuille-Michaud ^40^. *Salmonella enterica* Typhimurium SL1344 were procured from the American Type Culture Collection (Manassas, VA) and was maintained as previously described elsewhere ^37^. A single colony was inoculated into LB broth and grown for 8 h under aerobic conditions and then under oxygen-limiting conditions ^11,41^. Cells were infected with a multiplicity of infection (moi) of 1:10 for macrophages, 1:30 for EDMs and 1:100 for HEKs.

### Cell lines and cell culture

HEK293 cells were obtained from American Type Culture Collection (ATCC) and maintained in high glucose DMEM (Life Technologies) containing 10% fetal bovine serum and 100U/ml penicillin and streptomycin at 37°C in a 5% CO_2_ incubator. Control and ELMO1 small-hairpin RNA (shRNA) macrophage (J774) cells were maintained in high glucose Dulbecco’s modified Eagle’s medium (DMEM) containing 10% fetal calf serum, as described previously^6^. Cells were sub-cultured 24 hours prior to transfection. Transfections of plasmids were performed using Lipofectamine 2000 (Invitrogen) according to manufacturer’s protocol.

### shRNA lentiviral transduction

NOD2 MISSION shRNA Lentiviral Transduction Particles from Sigma Aldrich (TRCN0000066813) were used to stably down-regulate NOD2 in J774 macrophage. J774 cells were seeded in 96-well plates at 1.6x10^4^ cells per well for 24 hrs. Cell media supplemented with 8 ug/ml Polybrene (Sigma Aldrich, St Louis MO) was added to cells followed by 10ul of lentivirus particles (4.7 x 10^7^ VP/mL titre value). Next day, the media containing lentiviral particles were removed from wells and fresh media was added. For selection, 1mg/ml G418 (Cat# G8168, Sigma Aldrich) was added to cells 48hrs after lentivirus transduction. Cell media was changed every 2-3 days with fresh G418-containing media until resistant colonies were identified. RNA was extracted to determine knockdown efficacy and cell lines were kept under selective pressure using G418 containing media.

### Isolation of enteroids from the small intestine of the mouse

Intestinal crypts were isolated from the colonic tissue specimen by digesting with Collagenase type I [2 mg/ml; Invitrogen] and cultured in stem-cell enriched conditioned media with WNT 3a, R-spondin and Noggin as described before ^42–47^.

### The preparation of Enteroid-derived monolayers (EDMs)

To prepare EDMs, single cells from enteroids were plated in the presence of 5% conditioned media and diluted Matrigel (1:40) as done before ^47,48^. For all functional assays, experiments were performed multiple times with EDMs-derived from enteroids collected from at least 3 different mice, including both the genders.

### Isolation of Bone Marrow–Derived Macrophages

BMDMs were isolated following the protocol described elsewhere ^26,41^. Briefly, femur bones were collected from C57B6 mice after euthanization. The bone marrow cells were then flushed from the femur bones using 25 G needle and RPMI medium. The cells were then centrifuged, and the RBCs were lysed by incubating with 1X RBCs lysis buffer (Thermo Fisher Scientific) for 3 minutes. The remaining bone marrow cells were precipitated and resuspended in DMEM media containing 10% FBS, 20% LCCM (L929 cells conditioned media), and ciprofloxacin (10 µg/ml) and incubated at 37°C. After 3 days, the media was replaced with new media devoid of antibiotics.

### The measurement of Transepithelial electrical resistance (TEER)

TEER was measured in 24-well transwell plates following infection, using the STX2 electrodes with digital readout by EVOM2 (WPI) as described previously ^49^.

### Gentamicin protection assay

Approximately 3x10^5^ cells from WT, ELMO1^-/-^ and NOD2^-/-^ EDMs were plated onto a 0.4 µm pore trans well insert and infected with bacteria with moi 10. Bacterial internalization was determined after 12 h of infection of WT, ELMO1^-/-^ and NOD2^-/-^ EDMs with *AIEC-*LF82 following our previously published gentamicin protection assay ^6,9,10^. For transfected HEKs, approximately 4x10^5^ cells were plated 16h prior before infection and bacterial count was determined after 6h following infection. After infection cells were lysed with 1% TritonX-100, followed by serial dilution and plating on LB agar as done previously.

### Cytokine Assays

Supernatants were collected from the uninfected and infected cells, control or ELMO1/NOD2/both-knock down J774 cells after *AIEC*-LF82/ *Salmonella enterica* Typhimurium SL1344, infection and assessed for different cytokines such as IL-6, IL-1β, IL8, and MCP1 using the ELISA kit from R&D systems as per manufacturer’s protocol.

### RNA Preparation, Real-Time Reverse-Transcription Polymerase Chain Reaction

RNA isolation was performed using Zymol RNA extraction kit. cDNA was synthesized from the RNA using Quantinio cDNA synthesis kit (Quantabio) followed by quantitative RT-PCR using 2x SYBR Green qPCR Master Mix (Biotool™, USA) for target genes that was normalized to the endogenous control gene (18s rRNA) using the 2^−ΔΔCt^ method. Primers were designed using NCBI Primer Blast software. The primers used were as follows: 5’UTRNOD2 Forward 5’GGACCTGGACTCCTCCAAA3’ and Reverse 5’GCTGGGCTGAGAACACATAG3’.

### Expression Constructs

NOD2 mutant plasmids (HA-NOD2 R702W, HA-NOD2 G908R and HA-NOD2 L1007fs) were generated by site directed mutagenesis on HA-NOD2 plasmid (a gift from Dana Philpott) using the QuickChange II Site-Directed Mutagenesis kit (Agilent Technologies) according to manufacturer’s protocol. The CARD domain (amino acids 28–265) and LRR domain (amino acids 744–1040) of NOD2 were generated by PCR and cloned into pET-28a (+) plasmid vector (Novagene). All plasmid constructs and mutagenesis were verified by sequencing and protein expression was verified by western blot analysis.

### Immunoprecipitation and Western Blotting

Transfected HEK293 cells were lysed in NP buffer (1% NP-40, 0.5% Deoxycholate, 0.1% SDS in 1X PBS) with 1X Proteinase Inhibitor Cocktail added immediately before use. Whole cell lysate was centrifuged to separate proteins from cell debris and quantified using the Lowry assay. One mg of total protein lysate was incubated with 40ul of Ezview Red ANTI-FLAG M2 Affinity Gel (Cat # F2426-1ML, Sigma-Aldrich) overnight at 4°C. Beads were washed 4 times with NP lysis buffer at 1500 rpm for 2mins. Immunoprecipitates were eluted from the beads by resuspending beads in 50μl of 2X SDS-PAGE sample buffer and boiled for 10 mins at 100°C. Proteins were separated by running on a SDS-PAGE protein gel and then transferred onto Immobilon-P PVDF membrane. The membrane was immunoblotted with primary antibody followed by either Anti-mouse IgG HRP-linked Antibody (Cat # 7076S, Cell Signaling Tech) or Anti-rabbit IgG HRP-linked antibody (Cat #7074S, Cell Signaling Tech). Protein bands were detected using ECL (Amersham Biosciences).

### Purification of GST-NOD2-LRR

The bacterial colony (*Escherichia coli* BL21 expressing GST-NOD2-LRR) was picked from a freshly streaked plate, inoculated in 10 ml LB supplemented with 30 µg/ml ampicillin and incubated overnight at 37°C. The next day this 10 ml pre-culture was transferred to 1 L of LB supplemented with 30 µg/ml ampicillin and further incubated at 37°C (shaking at 220 rpm). After the OD of LB at A600 reaches to 0.6, isopropyl-β-D-thiogalactoside (IPTG) is added to a final concentration of 1 mM and incubated for overnight at 25°C. The cells were harvested by centrifugation for 30 min at 3750 rpm at 4°C. Cell pellet from 1L culture was resuspended in 10 ml of lysis buffer (50 mM Tris pH 8.8, 200 mM NaCl, 0.5 mM PMSF, 0.1% Tween-20, 0.2 % NP-40, 1 mM β-mercaptoethanol, 10% glycerol) and sonicated in 4°C cold room for 20 s four times, at 2 min intervals. The sonicated cell lysates were centrifuged for 30 min at 12000 rpm at 4°C. The supernatant was collected and incubated with GST beads for 1h at 4°C on rotator (GE Healthcare). GST beads were washed three times with phosphate buffered saline (PBS), and the attached proteins were eluted by adding reducing sample buffer. The purity of proteins was analyzed on 10% SDS-PAGE stained with Coomassie blue, quantified using BSA as standard, aliquoted and stored at -80°C.

### Pulldown protocol

Purified GST-NOD2-LRR was immobilized on GST beads for overnight at 4°C. The immobilized GST-NOD2-LRR was incubated with His-ELMO-CT and His-ELMO-FL proteins for binding for 4h at 4°C with constant tumbling in binding buffer (50 mM Tris–HCl (pH 7.4), 100 mM NaCl, 0.4% (v:v) Nonidet P-40, 10 mM MgCl2, 5 mM EDTA, 2 mM DTT, protease inhibitor mixture). Beads were washed four times with 1 ml of wash buffer (4.3 mM Na2HPO4, 1.4 mM KH2PO4 (pH 7.4), 137 mM NaCl, 2.7 mM KCl, 0.1% (v:v) Tween 20, 10 mM MgCl2, 5 mM EDTA, , 2 mM DTT) and boiled in Laemmli’s sample buffer. GST protein was used to detect nonspecific binding.

### Determination of intracellular ROS

Approximately 1 million cells were loaded with 5 μM dichlorofluorescin diacetate (H2DCFDA) according to standard procedures^50,51^. After incubation at 37°C for 15 min, the cells were washed and resuspended before being examined by flow cytometry (Guava® easyCyte Benchtop Flow Cytometer, Millipore). N-Acetyl l-cysteine (NAC) was used as scavenger of ROS at an optimized concentration of 10 mM as we have done before ^52^.

### Statistical analysis

Bacterial internalization, monocyte recruitment assays and ELISA results were expressed as the mean ± SEM and statistical significance was estimated using one-way ANOVA with Tukey’s test or a two-tailed Student’s t-test. Results were analyzed in the Graph pad Prism and considered significant if p values were < 0.05.

## Acknowledgment

We are grateful to Jasper Lee for his technical support in the organoid work. We are also thankful to Dr. Pradipta Ghosh for providing the access to equipment related to protein purification.

---

**Figure S1.**
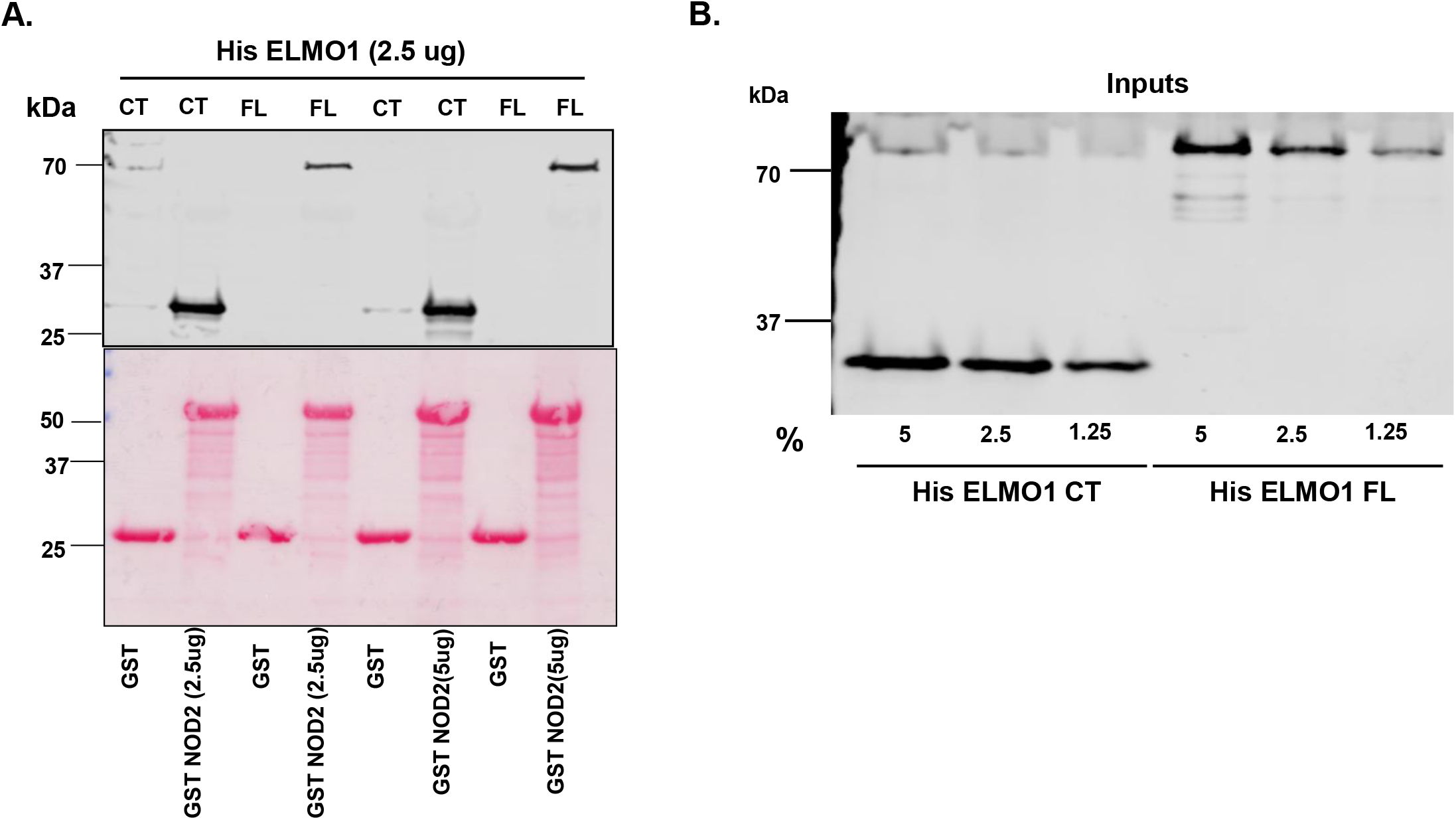
NOD2-LRR binds to C-terminus of ELMO1: **A.** GST-NOD2-LRR was immobilized on glutathione beads. Recombinant His-ELMO1-CT (∼3μg) and His-ELMO1-FL (∼3μg) was used in GST pulldown assays with GST or GST-NOD2-LRR. Bound His-ELMO1-CT or His-ELMO1-FL were visualized by immunoblotting using anti-His antibody. Equal loading of GST proteins was confirmed by Ponceau S staining in the lower panel. **B.** The input of His-ELMO1-CT and His-ELMO1-FL were shown in the right.

**Figure S2:**
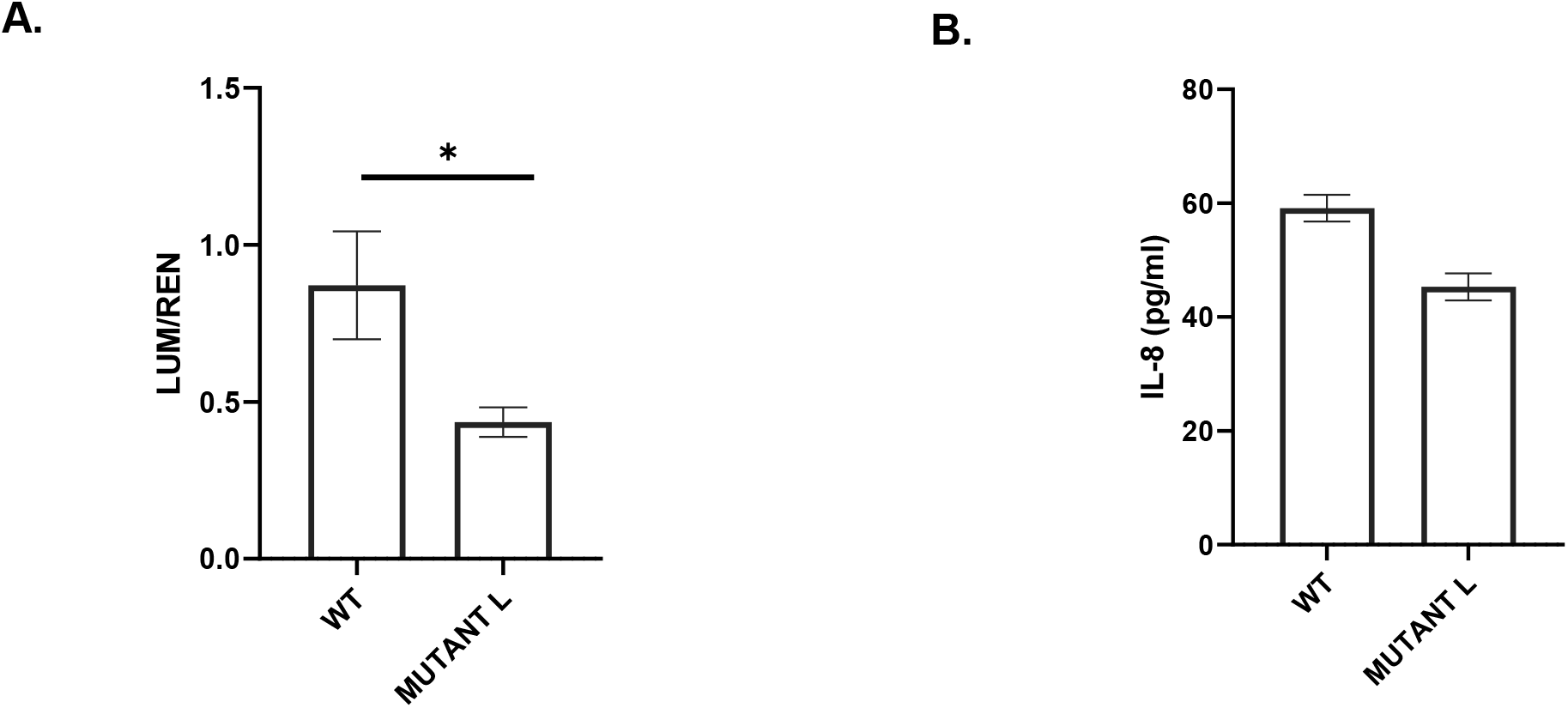
The NOD2 mutant affect NFkB activity followed by inflammatory cytokines. **A.** HEK293 cells were transfected with WT and NOD2L1007fs (Mutant L) followed by infection with *AIEC*-LF82 for 6h to perform the reporter assay. The relative luciferase activity was plotted. **B.** Supernatant was collected at 6h following MDP treatment and secreted IL-8 levels were measured by ELISA. Data displayed as mean ± SEM. Statistical significance was estimated using Mann Whitney test where **p < 0.01.

**Figure S3:**
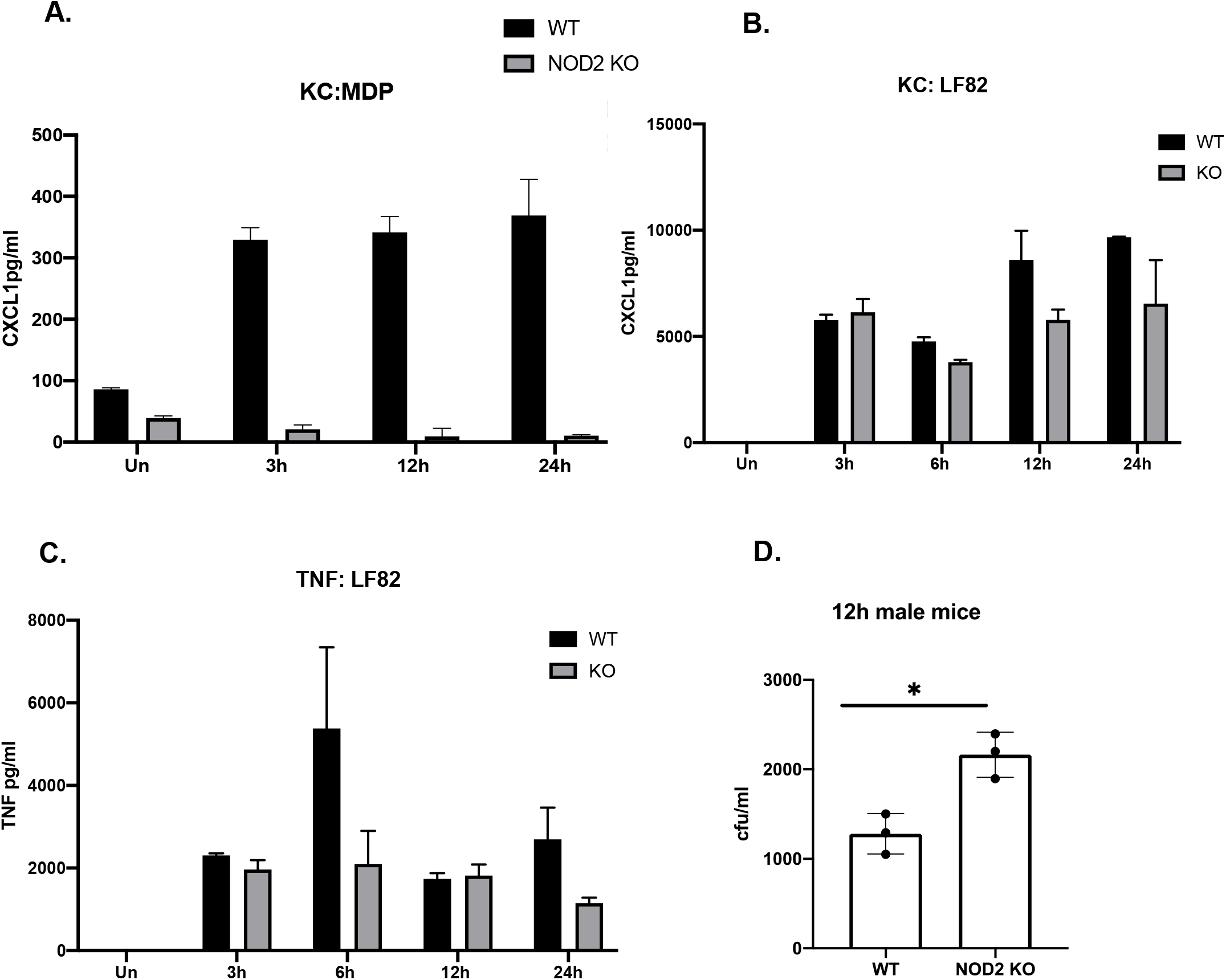
BMDMs were isolated from WT and Nod2 KO mice and either treated with MDP or infected with *AIEC*-LF82. Supernatant was collected at different time points. Graphs show concentration of (**A**) CXCL-1 upon MDP treatment. **B** CXCL-1 upon *AIEC*-LF82 infection **C**. TNF-a upon *AIEC*-LF82 infection**. D**. BMDMs were isolated from WT and Nod2 KO mice and infected with *AIEC*-LF82 for 1h and treated with high gentamicin for 90 min, and then left in low gentamicin. Graph shows number of bacterial colonies at 12hr.

**Figure S4:**
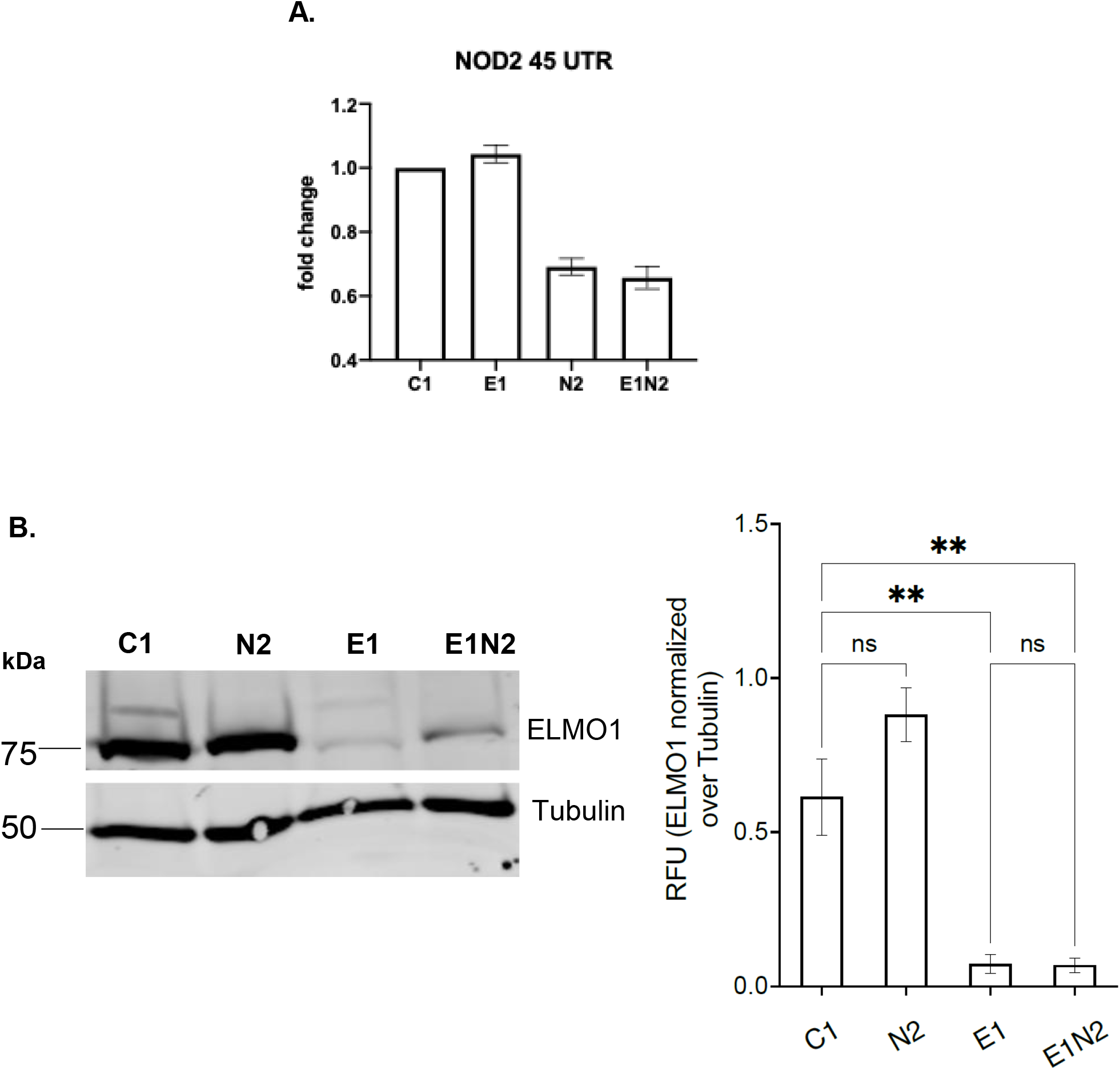
**A**. The expression of NOD2 was measured in C1, E1, N2 and E1N2 by qRT-PCR, **B**. Representative immunoblot of ELMO1 expression of in J774 cells control shRNA (C1), shELMO1 (E1), shNOD2 (N2) and shELMO1 & shNOD2 (E1N2). Densitometry was performed from 3 independent experiments. ELMO1 expression was normalized against tubulin expression and expressed as RFU values.

**Figure S5.**
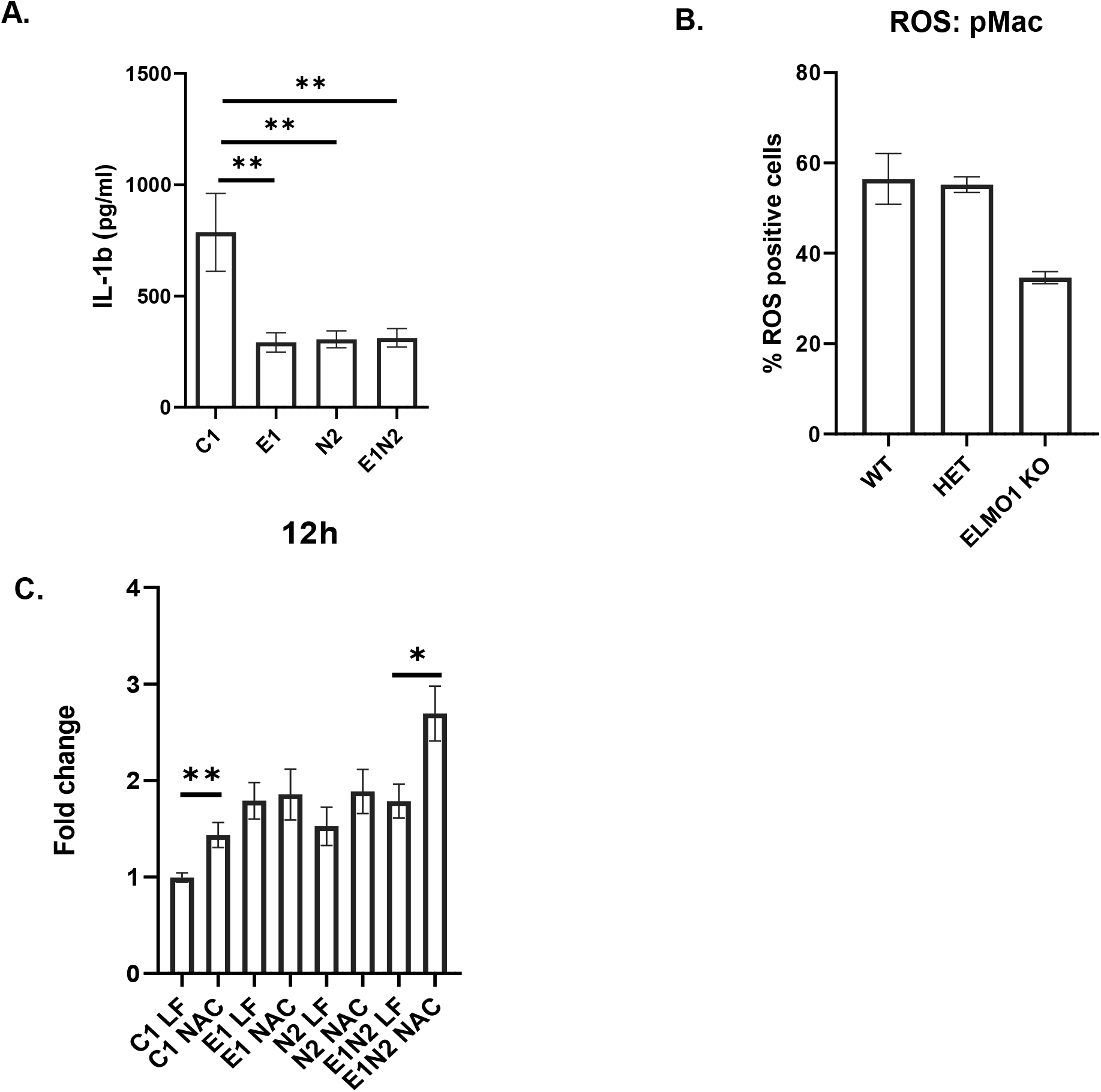
The impact of ELMO1-NOD2 interaction in immune response. **A.** J774 cells were infected with *Salmonella enterica* serovar Typhimurium strain SL1344 for 1h, after which extracellular bacteria were killed by gentamicin treatment. Cells were incubated for 12h Bar graphs display the concentration of pro-inflammatory cytokines (IL1β) released in supernatant at 12h after of *Salmonella* infection. **B.** Peritoneal macrophages were isolated from WT, Het and ELMO1 KO mice and infected with *AIEC*-LF82 for 1h and analyzed for ROS generation using flow cytometry. Graph shows percentage of cells expressing ROS. **C.** ROS Scavenger assay Data is displayed as mean ± SEM. Statistical significance was estimated using Mann Whitney test and one-way Anova *p < 0.05, **p < 0.01 and ***p < 0.001.

